# Complete neuroanatomy of dragonfly wings suggests direct sensing of aeroelastic deformations

**DOI:** 10.1101/2021.04.11.439336

**Authors:** Joseph Fabian, Igor Siwanowicz, Myriam Uhrhan, Masateru Maeda, Richard J Bomphrey, Huai-Ti Lin

**Affiliations:** Imperial College London; The University of Adelaide; HHMI Janelia Research Campus; Royal Veterinary College

## Abstract

Can mechanosensors in animal wings allow reconstruction of the wing aeroelastic states? Little is known about how flying animals utilize wing mechanosensation to monitor the dynamic state of their highly deformable wings. Odonata, dragonflies and damselflies, are a basal lineage of flying insects with excellent flight performance, and their wing mechanics have been studied extensively. Here, we present a comprehensive map of the wing sensory system for two Odonata species, including both the external sensor morphologies and internal neuroanatomy. We identified eight morphological classes of sensors; most were mechanosensors innervated by a single neuron. Their innervation patterns and morphologies minimize axon length and allow morphological latency compensation. We further mapped the major veins of another 13 Odonata species across 10 families and identified consistent sensor distribution patterns, with sensor count scaling with wing length. Finally, we constructed a high-fidelity finite element model of a dragonfly wing for structural analysis. Our dynamic loading simulations revealed features of the strain fields that wing sensor arrays could detect to encode different wing deformation states. Taken together, this work marks the first step toward an integrated understanding of fly-by-feel control in animal flight.

## Introduction

Animal locomotion relies on sensory feedback to implement reflexes (Clarac and Dando, 1973; Dickinson, 2000), plan future events (Mischiati et al., 2015; Lin and Leonardo, 2017) and accumulate experiences for motor learning (Singh and Scott, 2003). In terrestrial animals, proprioceptors in joints and muscles establish the timing and magnitude of important mechanical events during walking (Abelew et al., 2000). ‘Fly-by-feel’ describes how mechanosensory systems capture aerodynamic and inertial information to enable or enhance flight control. The wings of flying animals experience combined inertial-aerodynamic (‘aeroelastic’) loads, Coriolis forces, airflow stagnation and separation, and non-linear phenomena of vortex growth and shedding (Bomphrey and Godoy-Diana, 2018; Dickson et al., 2006; Combes and Daniel, 2003). The feather follicles of birds possess mechanosensors sensitive to feather motion under aeroelastic forces (Brown and Fedde, 1993; Carruthers et al., 2007), while the wings of bats and moths are populated with mechanosensors which are known to contribute to body stabilisation Sterbing-D’Angelo et al., 2011; Dickerson et al., 2014). The current literature on mechanical feedback during flight consists of isolated reports of specific sensor types, often accompanied by proposals of their functional significance for flight control. However, complete descriptions of all wing sensor arrangement have been elusive. Animal wings express a diverse array of mechanosensors each capturing a point measurement. As with sensory receptors in other organs, such as the eye, it is likely that arrays of receptors function in concert, each contributing a piece to a larger, time-dependent, sensory mosaic (Hardie, 1985; Samonds et al., 2018). Therefore, a comprehensive map of the mechanosensors found on wings is a critical first step towards understanding the function of sensory feedback during flight.

Fly-by-feel may be implemented differently in different flying animals, but the relatively simple nervous systems of insects simplifies the mapping process, and the absence of actuation beyond the wing base improves the tractability of mechanical analyses. Flapping flight relies on the phasic generation of precise aerodynamic forces through coordinated wing movements. Dragonflies are especially interesting due to their four independently controlled wings, each with adjustable amplitude, frequency, and angle of attack (Ruppel, 1989; Thomas, 2004). This enables a large wing state-space, each with unique, time-dependent, aerodynamic and inertial characteristics. Dragonflies can fly by synchronised flapping of all four wings, out-of-phase flapping of forewings and hindwings, flapping of the hindwings only, gliding, and even in mechanically-linked tandem mating flights. Practically, dragonflies’ exposed wing veins and transparent, scaleless membranes, facilitate the characterization of external and internal anatomy.

Sensory feedback from insect wings has been studied by neuroscientists and engineers since the early 1900s (Vogel, 1911; Lehr, 1914; Gettrup, 1966; Wilson, 1961) following the discovery of mechanosensors on butterfly wings. Further electrophysiological studies described afferent signals from mechanosensors called campaniform sensilla in fly wings (Dickinson, 1990). These neurons tend to fire a single action potential per wing stroke, so they cannot provide continuous strain estimation. Instead, strain is believed to be encoded predominantly through spike timing, which varies with the neurons spiking strain threshold, and the temporal dynamics of local strain Dickerson et al., 2014; Yarger and Fox, 2018). Unlike most other insects, flies have specialized club-like structures called halteres, derived from their reduced hindwings. Instead of producing aerodynamic forces during flight, the halteres express strain sensors which allow estimation of the insect’s body rotation through the detection of the Coriolis forces (Pringle, 1948). All flapping insect wings experience Coriolis forces, and recent work has demonstrated that strain sensor arrays on wings also function as inertial sensors, much like the halteres (Pratt et al., 2017; Jankauski et al., 2017; Eberle et al., 2015). In fact, to detect body rotation through a flapping wing, only relatively few sparsely distributed strain sensors are needed (Hinson and Morgansen, 2015). This is consistent with the sparse distribution of campaniform sensilla found in some insects (Dickinson and Palka, 1987). However, the wing-mounted sensors are also well-positioned to monitor the instantaneous aeroelastic wing deformation and local flow velocities on the wings themselves. This line of investigation is far less developed and can impact our understanding of flight control in flying insects, flying vertebrates and also inspire novel, engineered flight controllers.

To pave the way for comprehensive and mechanistic studies of animal fly-by-feel control, we present the most complete description of an animal wing mechanosensory system to date. In the process, we answer several key questions about the neuronal architecture and biomechanics of this system. Specifically, we show the neural innervation, locations and morphologies of all sensor types found on the wings of Odonata. We discuss these findings in the context of sensory performance, metabolic costs, wing damage and redundancy. Finally, we present the most anatomically accurate finite element model of a dragonfly wing blade to demonstrate the discrete strain fields that propagate through the structure.

## Results

### 1. Within an extensive network, sensory neurons follow direct paths to mechanosensors in Odonata wings

#### Which regions of the wing blade are innervated by sensory neurons?

To visualise the neuronal innervation within Odonata wings, we imaged the complete collection of wing neurons and associated cuticular structures via confocal microscopy (see Methods). For comparison, we visualised wings from a small dragonfly species *Perithemis tenera* and a similar-sized damselfly *Argia apicalis*. Some of these neurons’ axons run the entire wing length, making them among the longest in insects. All axons follow a relatively direct path and terminate with a soma which is often located directly under an external cuticular structure (Figure 1A,B). Unlike the leading edge and the longitudinal veins, the trailing edge of *P. tenera* wings has no dedicated axon tract. Instead, it is innervated locally by extensions of the closest longitudinal vein tracts. The venation of damselfly wings consists of well-aligned rectangular cells, resembling a modern city grid. The innervation pattern of the damselfly (*A. apicalis*) wing is similar to that of the dragonfly (*P. tenera*), with some exceptions (Figure 1C). Unlike the dragonfly wing, the trailing edge of the damselfly wing is innervated directly, with a tract of axons running along the length of the trailing edge.

**Figure 1.**
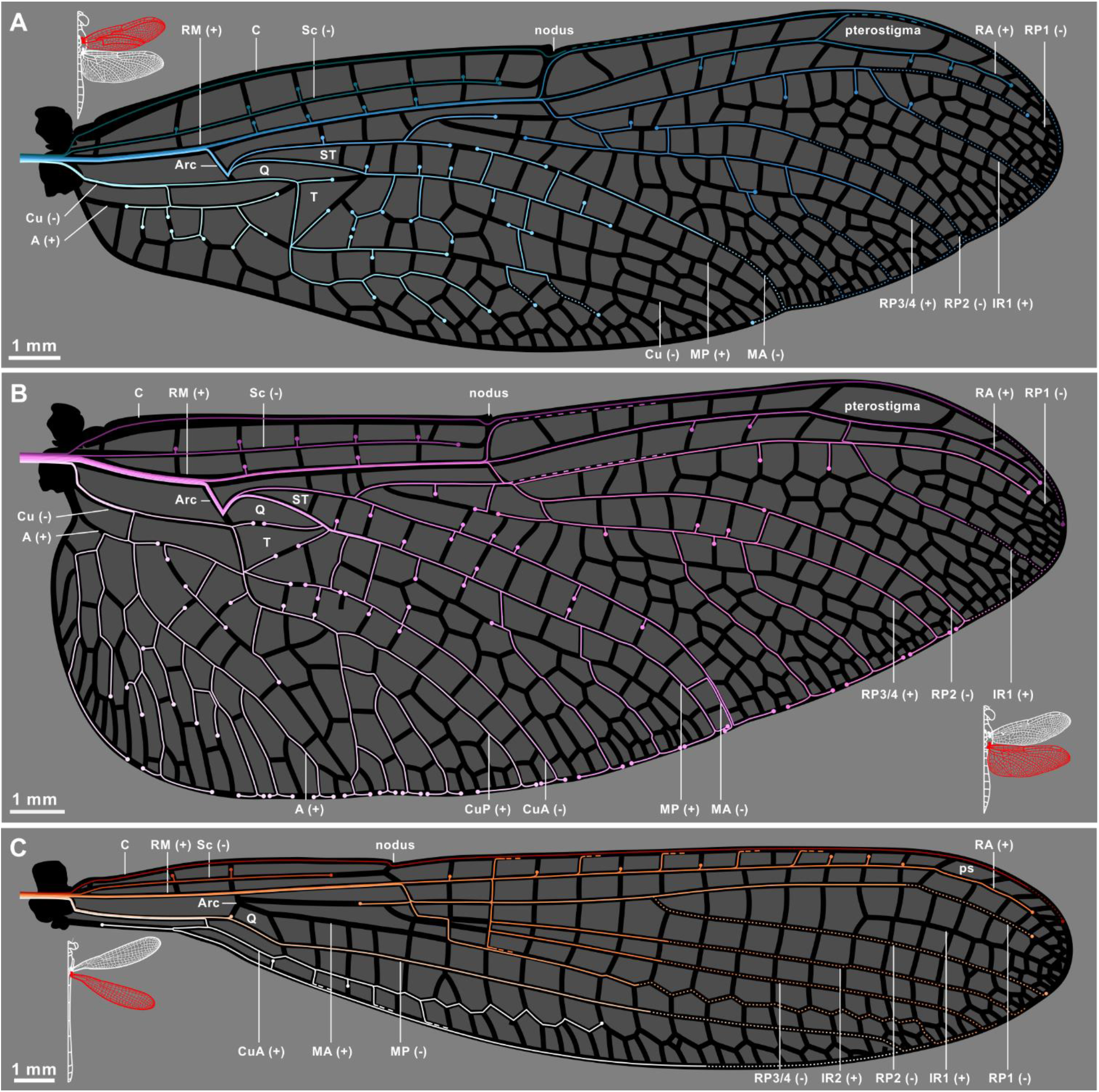
The axon routing pattern of selected Odonates. Neuronal innervation diagram of male Eastern Amberwing dragonfly (*Perithemis tenera*) forwing (A), hindwing (B) and a male Blue Fronted Dancer damselfly (*Argia apicalis*) hindwing (C). A dedicated afference wing nerve branches out into different veins from the wing base. The color tones of axonal tracks were chosen arbitrarily for readability. They do not imply hierarchy or relationship of the tracks. Dots indicate terminal mechanosensory cell body for each track; dashed lines denote merging tracks; dotted lines represent tracks that were interpolated from incomplete back-fill images and mechanosensors found on the veins. Major veins and structural elements of the wings are labeled following the system of Riek and Kukalova-Peck (1984), Bechly (1995) and Trueman and Rowe (2019): C-costa; Sc - subcosta; RM - radius/media; Cu - cubitus; A - anal; Arc - arculus; T - triangle; ST - supratriangle; Q - quadrilateral. The corrugations of the wings/ topology of veins are represented by “+” for “ridges” (dorsally convex) and “−” for “valleys” (dorsally concave).

### 2. Variation and optimisation of axon routing in the wing

#### How stereotyped is the axonal routing and what type of variations exist?

The complex venation pattern of dragonfly wings allows many different axon pathing options. We compared the routes axons take in a set of male and female *P. tenera* wings and highlight regions where we observed differences in axon routing (Figure 2A). We performed a larger survey of wing routing through a key feature of Odonata wings: the unidirectional flexion joint (i.e. nodus). The nodus contains a significant amount of elastic resilin, a material which allows larger flexion and elastic energy storage during wing motion (Rajabi et al., 2017; Yazawa et al., 2018). Passing an axon bundle across an articulating joint is more structurally complex than a fixed vein. Interestingly, we see that in some individuals axon bundles from the proximal costa pass through the nodus joint to innervate the distal costa, whereas in other individuals the distal costa is innervated by branches of subcosta bundles, by-passing the nodus (Fig 2B).

**Figure 2.**
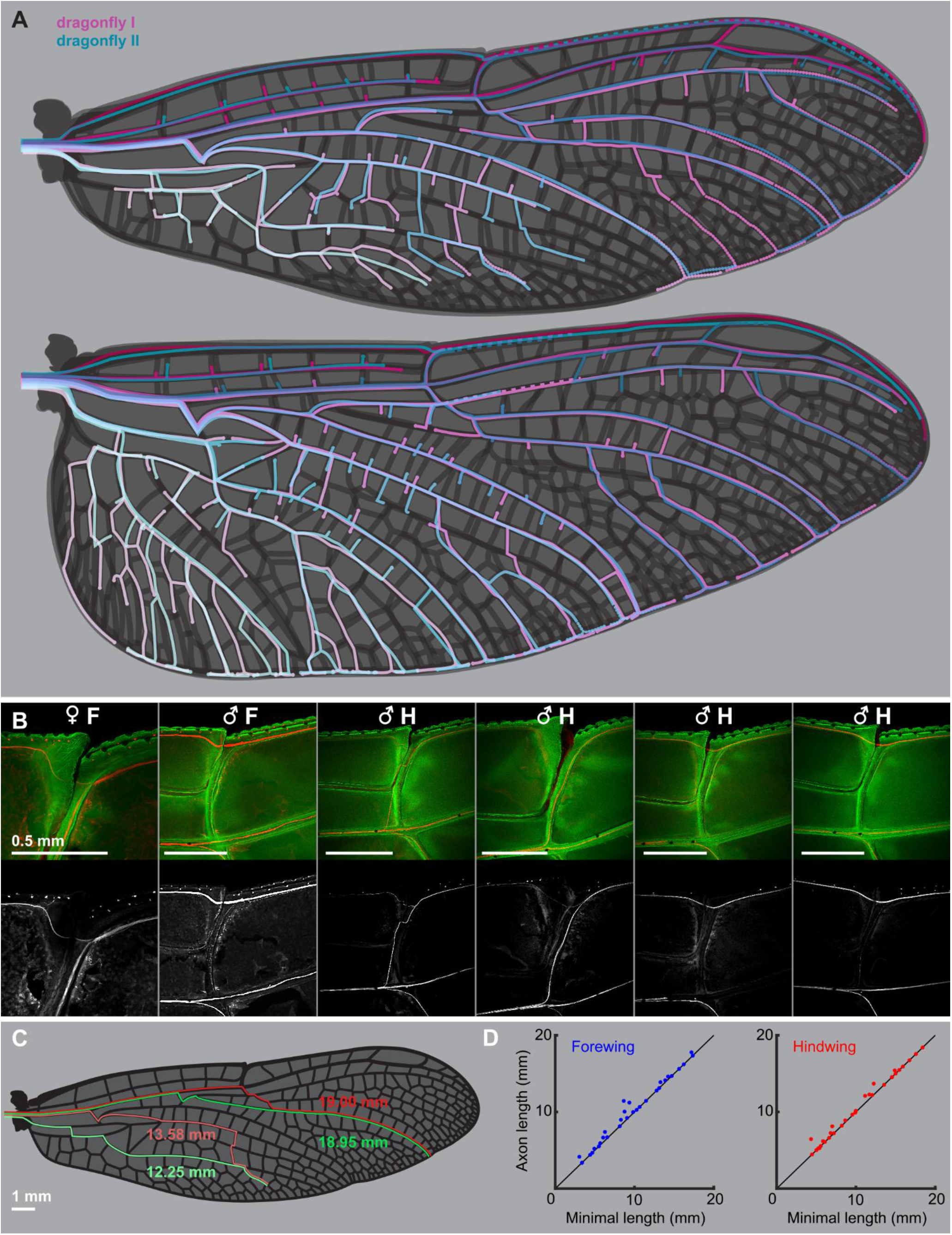
(A) Variability of axonal tracks in two dragonflies (*Perithemis tenera*) fore- and hindwings. The dragonfly II wings were superimposed on the dragonfly I’s, scaled, and warped in Adobe illustrator to align the wing margins and main veins. Each vein carries a single nerve. (B) Variability of the axonal track passage at the nodus of *P. tenera*: axons of the leading edge wing margin sensors distal to nodus may continue down the costa or take an alternative path down the radius vein. The path does not seem to depend on the sex of the insect and the variability occurs in both fore (F) and hind (H) wings. (C) Real paths (red) and minimal distance paths (green) of axons innervating a wing margin bristle bump complex neuron (anterior pair) and a campaniform sensillum (posterior pair) in a male (dragonfly I) forewing. (D) The real path distance and minimal path distance for 31 pseudorandomly selected sensors on the forewing and hindwing of a dragonfly wing.

We hypothesise that axon length minimisation is a primary factor determining the organisation of axon paths in the dragonfly wing. To test this we pseudorandomly selected 31 evenly distributed sensors on the forewing and hindwing of one *P. tenera* sample and compared the true axon length for the selected path against the shortest feasible path from sensor to wing hinge (Figure 2C–D). Combining data from both wings, the total axon length to innervate the 62 selected sensors was only 3.3% longer than the shortest possible routing solution, with the axons of 53.2% of sensors taking the shortest viable path. Sensors which took suboptimal routes for axon length minimisation tended to make common “mistakes”, particularly around the arculus and nodus, regions that play important structural roles in flapping flight (Wootton, 1992). Our data suggests that the organisation of axons in the dragonfly wing is well optimised to reduce total wiring length, within the constraints of the mechanical roles of different wing veins.

### 3. Wing sensor morphologies and classification

#### What are the sensors found on an Odonata wing?

Odonata wing membranes are smooth, but their veins are covered in microscopic structures with different roles (Rajabi et al., 2011; Gao et al., 2013). To identify sensory structures, we combine imaging data of external morphologies and internal anatomy. Based on our current understanding of insect wing sensors, we classified the wing sensory structures into eight classes: wing margin bristle-bump complexes (Figure 3A,B,C); bristle-bump complexes (Figure 3D); isolated bristles of varying length (Figure 3E,F); campaniform sensilla with associated structures (Figure 3G,H,J); isolated campaniform sensilla (Figure 3I,K,L); campaniform sensilla fields (Figure 3M,N); hair plates (Figure 3N,O); and multipolar cells (Figure 3P,Q,R).

**Figure 3.**
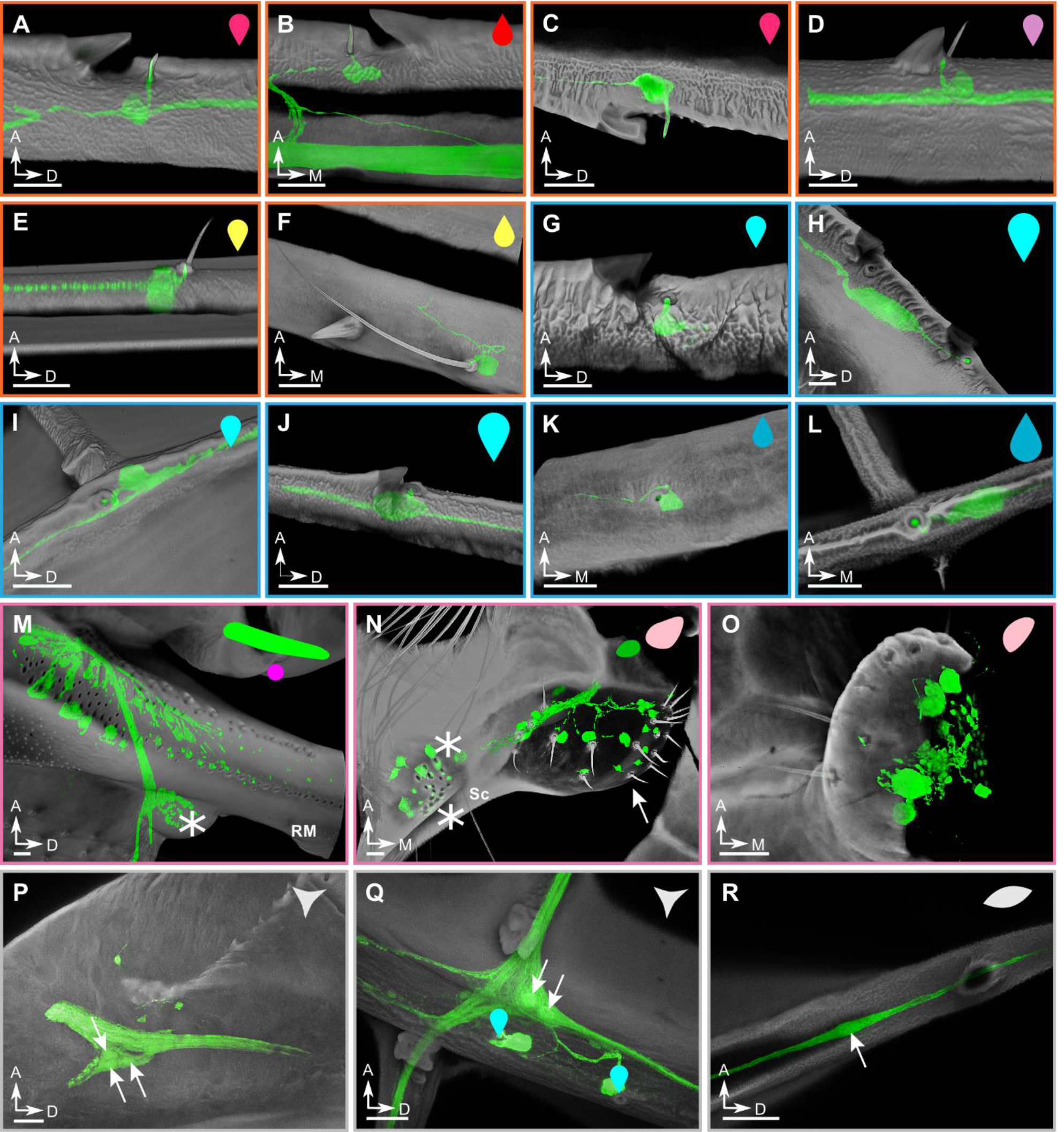
The classification and morphology of all wing sensors. Examples of sensors found on the wings of the dragonfly *Perithemis tenera*. The scale bars are all 25 μm. (A-F) Candidate airflow sensors: (A) dorsal costa margin bristle-bump; (B) ventral costa margin bristle-bump; (C) trailing edge margin bristle-bumps. All three examples show the double-innervated type of a bristle which consist only 25% of all the wing margin bristles. All other bristles have only one dendrite innervating the bristle base (not shown). (D) radius bristle-bump; all bristles of this type are innervated by one neuron at the base. (E) Short bristle of the type present on several major veins. (F) Long bristle of the type present on the medial part of major veins. (G-L) Strain sensors: (G) dorsal costa campaniform sensillum (CS); (H) large dorsal costa CS; (I) dorsal cross vein CS; (J) large radius anterior CS immediately distal to pterostigma; (K) ventral subcosta CS; (L) terminal ventral CS of the radius posterior 1 (RP1) vein. (M-O) Wing base sensory fields: (M) crevice organ; two parallel fields of directionally tuned elliptical CS at the base of radius/media (RM) vein. Asterisk marks the dorsal insertion site of a wing base chordotonal organ; (N) hair plate (arrow) and two adjacent CS fields (asterisks) ventrally at the base of subcosta; (O) hair plate ventrally at the base of cubitus. (P-R) Multipolar and bipolar receptors: (P) multipolar receptor at the base of costa; arrows point to/indicate cell bodies; (Q) multipolar receptor located at the junction of the anal vein and the first/medial cross vein connecting it to cubitus. Two adjacent dorsal CS are indicated with cyan markers. (R) Bipolar receptor. All example images are from the right forewing, with cuticle in grey, and neurons in green. The symbols in the upper right corner of each panel are notations to show sensor type and dorsal/ventral placement which will be used in Fig 4 and Fig 5.

**Figure 4.**
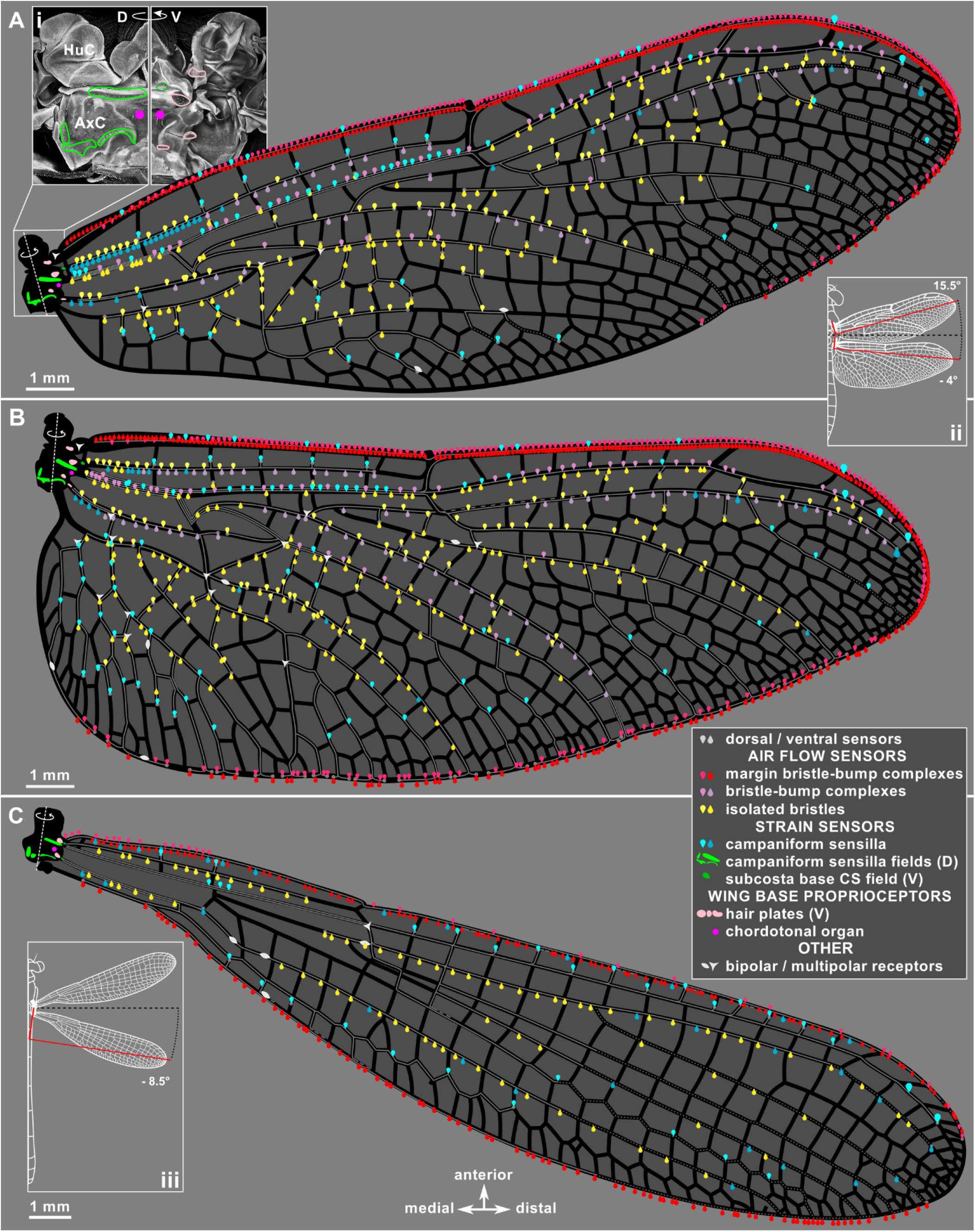
A sensory map of Odonate wings. Distribution of all confirmed sensors on the *Perithemis tenera* dragonfly fore-(A) and hind (B) wings and a hindwing of *Argia apicalis* damselfly (C). All sensor notations follow Fig 3 and the figure legend. Inset (i): maximum intensity projection showing *P. tenera* right forewing base dorsally and ventrally with the sensor fields outlined. Insets (ii) and (iii): diagrams showing the wings’ natural resting sweep angles; the red reference lines mark the wing span axis perpendicular to the anatomical wing hinges.

**Figure 5.**
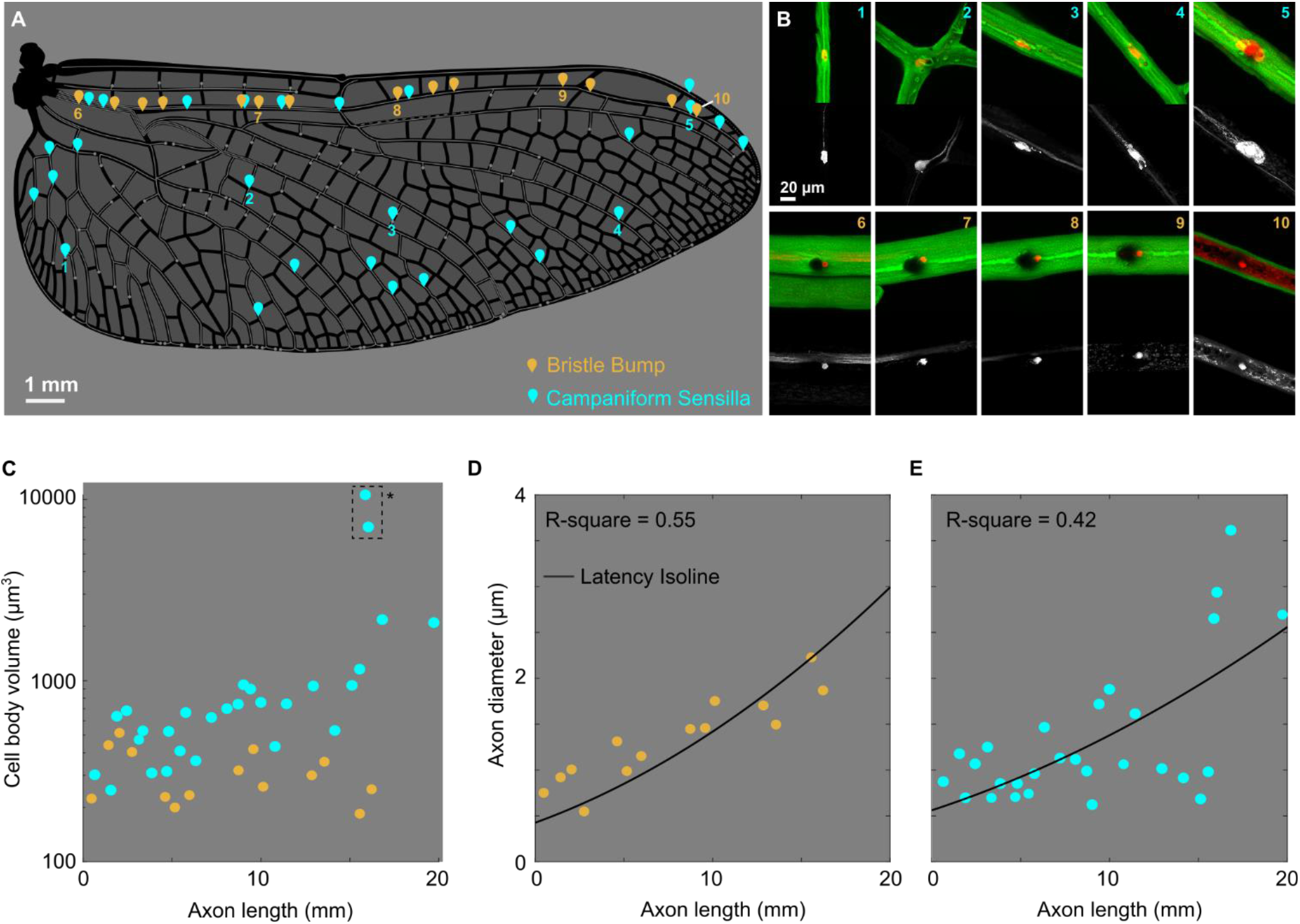
Cell body volume and axon diameter quantification. (A) Positions of selected wing sensors on the right hindwing of *Perithemis tenera*. Cyan - campaniform sensilla (CS) orange - bristle bump complexes. (B) Images of the cell body and axon of wing sensors, with cuticle in green and neurons in red. Greyscale images underneath show the neuron channel alone. (C) Cell body volume for wing sensors outlined in A. (D-E) Axon diameter for bristle bumps (D) and campaniform sensilla (E), black line indicates the axon diameter required to achieve equal response latency for different axon lengths.

The costa vein (leading edge) and trailing edge of the wing are densely populated with sensory bristles (~30 μm long in *P. tenera*), associated with dorsal or ventral serrations (Figure 3A–C). Similar bristles have been found on the wing margin of moths, sensitive to directional vibratory airflow (Yoshida and Emoto, 2011), and in Drosophila, where stimulation triggers a defensive leg kicking response (Li et al., 2016). In the dragonfly, each bristle attaches to the dendrite of a sensory neuron at its base, and in up to 25% of cases a second sensory neuron sends its dendrite through the shaft of the bristle to its tip. These are the only external wing sensors that are innervated by more than one neuron in Odonata.

The thickest longitudinal veins (subcosta, radius, cubitus, media) contain short (10–15 μm) bristles, each paired with a small saw-tooth shaped bump (Figure 3D). The shape of these paired bumps is highly stereotyped across the wing, and across multiple species, as is the spatial relationship between the bump and bristle. Each bristle lies several micrometres distal from the bump, which protrudes from the vein at a shallow gradient on its medial side, and a steep gradient on its distal side.

A variety of strain sensitive campaniform sensilla are found on the costa (Figure 3G,H), and several other longitudinal veins (Figure 3I–L), often paired with the same type of saw-toothed cuticular bumps described for short bristles. We observe isolated campaniform sensilla on many of the joints connecting the longitudinal radius vein to cross veins (Figure 3L). At the base of the radius, we see five fields of campaniform sensilla, with each sensillum innervated by an associated sensory neuron (Figure 3M). These sensor fields match the crevice organs described by Simmons (Simmons, 1978), sensors whose name at the time reflected their resemblance of small pits on the cuticle. Fields of campaniforms with high aspect ratio pits are associated with directional strain sensing (Spinola and Chapman, 1975; Vincent et al., 2007; Zill and Moran, 1981). Specifically, two such fields on the subcosta have an almost orthogonal directional bias which is conserved across odonates. We also identify several hair plates on the wing base (Figure 3N,O), as well as a chordotonal organ, identified by its characteristic anatomy (Field and Matheson, 1998; Wolfrum, 1990) (Figure 3M). These sensory structures are typically associated with proprioception in insects (Kutsch et al, 1980; Burrows, 1996). Finally, we observe multiple multipolar neurons located at some joints between wing veins (Figure 3Q,R) with unknown function and no associated cuticular structure.

### 4. A complete sensor map of Odonata wings

#### How many sensors are on the wing and where are they located?

Excluding the wing-base campaniform fields, we found a total of 771 sensors on the forewing and 894 sensors on the hindwing of a dragonfly (*Perithemis tenera*), and 358 sensors on the hindwing of a damselfly (*Argia apicalis*). Extrapolating from this, we estimate this small dragonfly has well over 3000 wing sensors on its four wings, and the damselfly approximately half as many. We found a total of 84 and 98 isolated campaniform sensilla on fore- and hindwing of *Perithemis tenera*, respectively, with a bias towards the dorsal surface. The abundance of each sensor type varies considerably, and bristles significantly outweigh campaniform sensilla on most veins. Interestingly, the damselfly wings studied are completely devoid of bristle-bump complexes or dorsal bristles (Figure 7C).

### 5. Neuron size and axon length

#### Is there any morphological adaptation to account for the sensory latency of wing sensory neurons?

The dragonfly wing is a long structure, and the axon lengths between sensors on proximal or distal regions of the wing (Figure 5A,B) could introduce significant temporal differences in spike arrival at the ganglia. While such difference can be accounted for in the sensory representation scheme, a better solution is to remove the temporal difference entirely through anatomical modifications. Specifically, distal sensors can compensate for their longer axons by increasing conduction speed via enlarged axon diameters. We measured cell body volume and the axon diameter for sensors at different span-wise locations on the wing. Cell body volume is approximately constant for bristle bumps, but tended to scale with axon length for campaniforms (figure 5C), with two abnormally large cells located near the wing tip, which we refer to as “giant wing tip campaniforms”, highlighted by a dashed line. We found that axon diameters tend to grow as you move along the wing span, both for bristle bumps (figure 5C) and campaniform sensilla (figure 5D). Based on the cable theory for neural conduction speed (Kandel et al, 2000; asaki, 2004), we derived a relationship between the axon diameter and length for “isochronic” scaling (see Methods). The axon diameter of bristle bumps and campaniform sensilla resembles values predicted by our latency isoline (R^2^ = 0.55 and R^2^ = 0.42 respectively), suggesting most wing sensors scale their axon diameters to maintain a similar sensory latency across the wing.

### 6. Giant campaniform sensilla near the pterostigma

#### How are giant wing tip campaniform sensilla distributed across Odonata species?

We systematically searched for campaniform distribution patterns near the wing tip across 15 Odonata species (Fig 6). All but one dragonfly species studied (*Anax junius*) have an isolated campaniform sensillum immediately distal to the pterostigma on the costa and radius veins, but they were absent in all three damselfly species. In *P. tenera* where we performed internal neuron size measurements, these two giant campaniform sensilla can be seen in Figure 5. The cell bodies are so large that we observe a significant bulge in radius vein to accommodate them (Sup Fig 2F).

**Figure 6.**
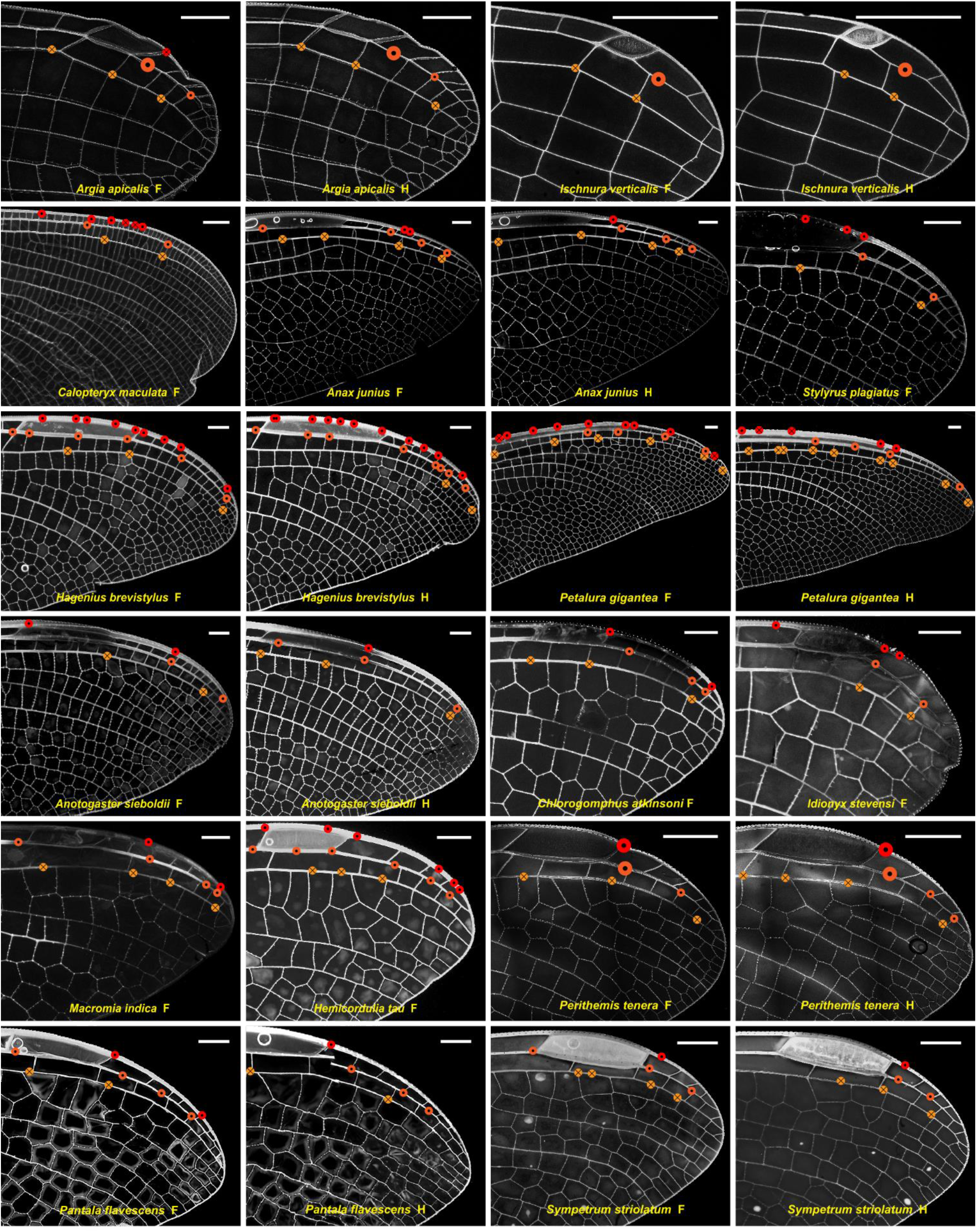
Positions of costa (red), radius anterior (dark orange) and radius posterior (light orange) campaniform sensilla at the wing tips of studied odonates. Dots denote dorsal, crosses – ventral sensors. Enlarged markers indicate campaniform sensilla innervated by sensory neurons possessing significantly larger soma as seen in backfills for *Argia apicalis* and *Perithemis tenera*, and inferred from a prominent bulge on the vein in *Ischnura verticalis*, likely expanded to accommodate a large cell body. Scale bars: 1 mm.

### 7. Sensor distribution across Odonata species

#### Do sensor densities inform some consistent patterns across odonate species?

Odonata is a diverse insect order, phylogenetically close to the first flying insects, with wingspans, wingbeat frequencies and flight speeds varying significantly across species (Bomphrey et al., 2016). How consistent are the classes and distribution of wing mechanosensors across different species of Odonata? We imaged the external morphology of wing sensors in 15 species (12 dragonflies and three damselflies) from 10 of the most studied families of odonates. Our analyses focused on three major longitudinal veins near the leading edge: the costa, subcosta and radius, as they are anatomically similar across all species, express a diverse range of sensors, cover most or all of the wingspan, and bear most of the load during flight (Büsse and Hörnschemeyer, 2013; Wooton, 1992).

All species studied maintain the same relationship between wing corrugation and sensor class (Figure 7A–H). Bristle-bump complexes and campaniform sensilla are found on the ridges (ventral side of subcosta and dorsal side of radius) and isolated bristles in the valleys (dorsal side subcosta and ventral side radius). Margin bristle-bump complexes and wing base campaniform fields are found in all 15 species (Figure 7A), but bristle-bump complexes (Figure 7C,F), campaniform sensilla (Figure 7B,D,G), and isolated bristles (Figure 7E,H) are missing in some species. The differences across species are most evident on the subcosta, where only dragonfly species express bristle-bump complexes and isolated bristles (Figure 7C,E).

**Figure 7.**
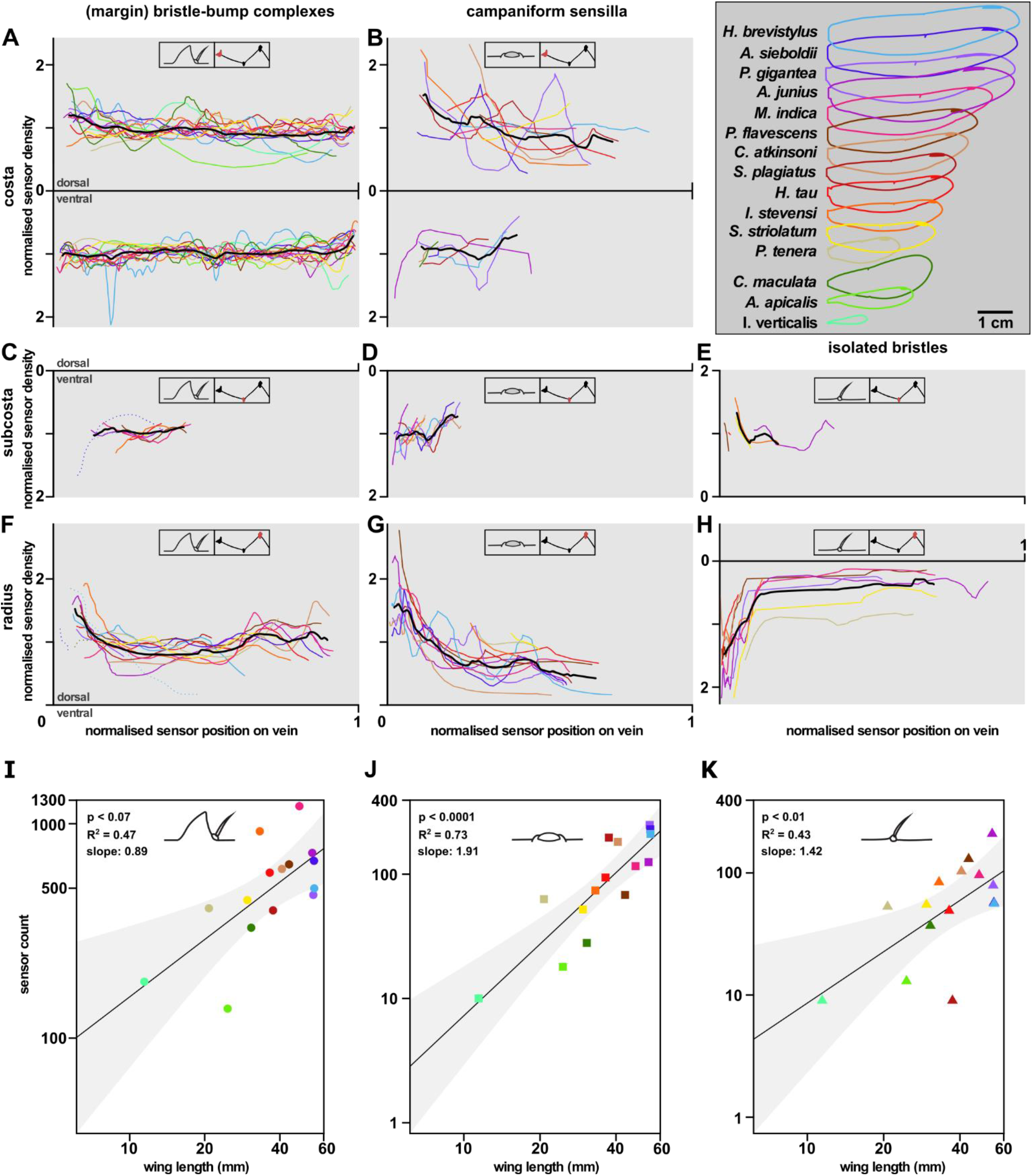
A comparison of sensor distribution across dragonfly families. Sensor density distribution (A-H) and sensor count (I-K) of the wing sensor types (bristle-bump complexes, campaniform sensilla and isolated bristles) across 15 Odonate species. Normalised sensor density is shown for dorsal and ventral side over normalised sensor position on three longitudinal veins, costa (A-B), subcosta (C-E) and radius (F-H). The insets in each panel highlight sensor type and the respective vein. Sensor density is normalized by the mean of each density curve. Sensor position is normalised by the length of the respective right forewing. The black bold line shows the mean of all species in each plot. (I-K) Sensor count is the sum for each sensor type of all three longitudinal veins. The sensor count is plotted over the respective wing length for each species. Black line shows the ordinary least squares (OLS) regression line and the grey shaded area shows the 95% confidence interval. Each data point is color coded by the respective species (see legend in upper right panel).

On the costa, the normalised density of wing margin bristles is relatively constant across the wing length for all species (Figure 7A). However, the damselflies *I. verticalis*, *C. maculata* and the dragonflies *P. tenera*, *S. plagiatus* and *I. stevensi* show local density peaks near the nodus (approximately half wing length) on the dorsal side. For all species but *P. gigantea*, sensor densities are higher on the ventral side compared to the dorsal (Table S1). Smaller wings have higher wing margin bristle-bump sensor density, with *I. verticalis*, *P. tenera* and *I. stevensi* having an average density of 7 sensors/mm, 8 sensors/mm and 10 sensors/mm respectively compared to 4 sensors/mm average density for *A. sieboldii* and *H. brevistylus* on the dorsal side of the costa. The density of campaniform sensilla along the major veins decreases gradually from wing hinge to tip (Figure 7B,D,G).

The cross-species comparison reveals that sensor counts increase with wing length for all three sensor types (Figure 7I–K). Given the data points available, bristle-bump complex (OLS slope 0.89; 95% CI [0.29, 1.5] and isolated bristle (OLS slope 1.42; 95% CI [0.39, 2.45]) counts resemble isometric scaling with wing length, while the campaniform sensilla counts scale faster than the isometric line (OLS slope 1.91; 95% CI [1.18, 2.6]).

### 8. The strain distribution and venation patterns

#### How does the wing venation impact mechanical strain distribution and therefore the strain sensory encoding?

Insect wings are lightweight but sufficiently stiff to support aerodynamic and inertial loads. Their intricate venation and extremely thin membrane (~10 μm thickness) make them challenging to image and model. As a result, insect wings are often simplified to thin plates with uniform thickness for aerodynamic or structural modelling (Liu et al., 2016). In reality, strain fields on the wing do not spread uniformly. Instead, they are channelled through the veins which also happen to house all the mechanosensors. To visualize how the venation pattern impacts strain fields during flapping, we created the most anatomically accurate dragonfly wing model to date for simple deformation analyses.

We constructed a finite element wing model from μCT data (see Methods) for a *S. striolatum* dragonfly hindwing. To understand the simplest dynamic loading behaviour of the structure, we subjected our model wing to a sinusoidal flapping oscillation around a single axis (parallel to the wing plane and roughly perpendicular to the spanwise axis) in a vacuum to observe strain field propagation given the venation pattern. The surface spanwise strain contours of four snapshots (Figure 8A–D) show a high concentration of strain at the longitudinal veins, especially near the wing base. The spanwise strain is greatest at the stroke reversals (Figure 8A,C); the proximal portion of the radius experiences compression (negative strain, in blue) at the transition from upstroke to downstroke (Figure 8A), while the same vein experiences tension (positive strain, in red) at the transition from downstroke to upstroke (Figure 8C). On the other hand, the proximal subcosta and cubitus (two longitudinal veins adjacent to the radius) have the opposite sign. This is because the corrugation offsets some longitudinal veins away from the neutral plane of bending. We also quantified the spanwise strain along two major longitudinal veins (subcosta and radius, Figure 8E–J) where we found campaniform sensilla on their corresponding ‘ridge’ sides (ventral side for subcostal and dorsal side for radius). Each of these veins shows a consistent spatial distribution of strain along their length (Figure 8G,H), but with a large temporal variation in magnitude over the flapping cycle (Figure 8I,J).

**Figure 8.**
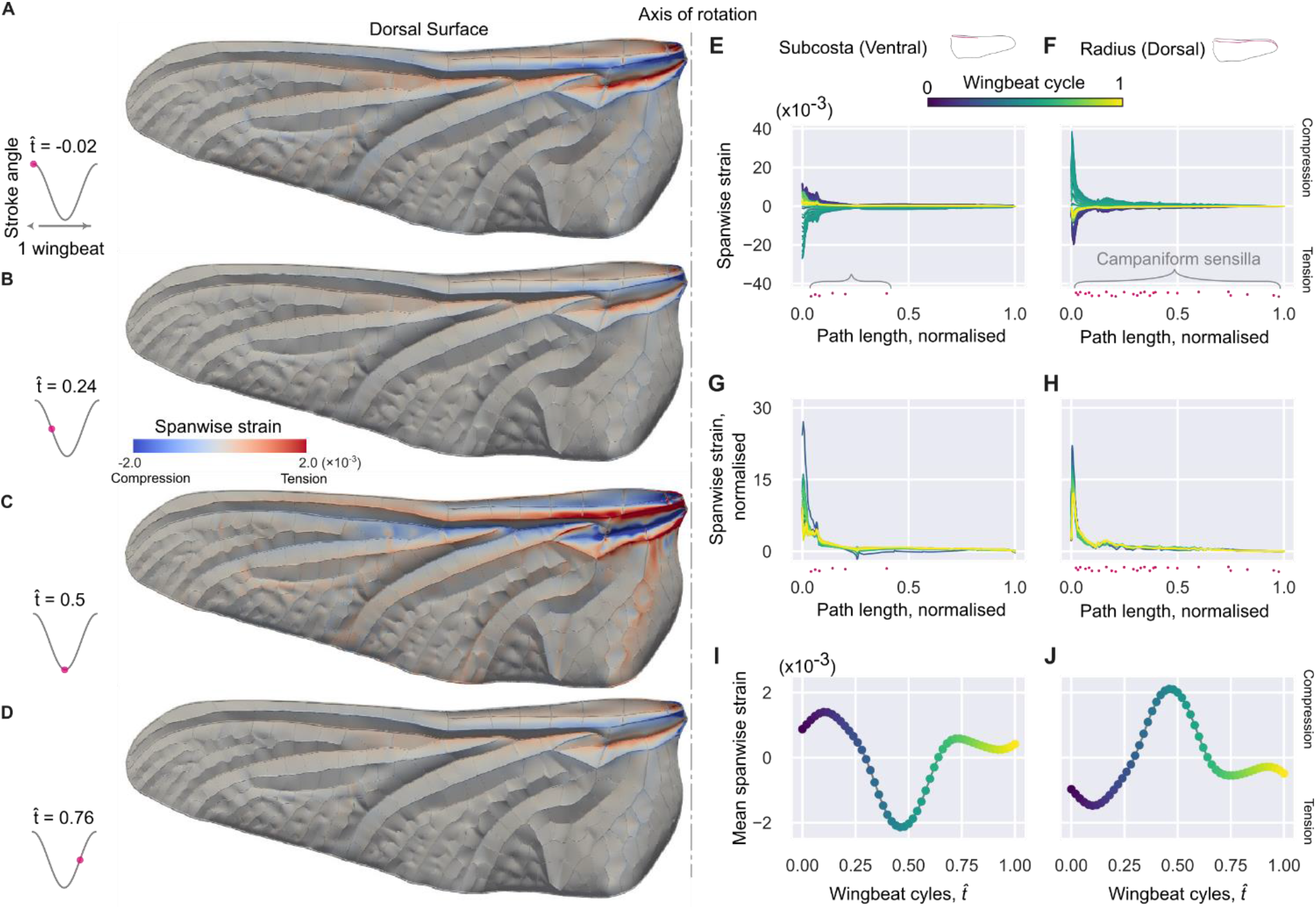
Strain distributions on a simulated flapping wing. (A–D) Spanwise strain (spanwise component of normal elastic strain) contour over the wing surface for four time instances in a wing stroke cycle: Beginning of downstroke (A), mid-downstroke (B), beginning of upstroke (C), and mid-upstroke (D). A hindwing of a Sympetrum striolatum was reconstructed in 3D for this simulation. (E,F) Spanwise strain along ventral subcosta and dorsal radius where campaniform sensilla can be found. The colour map represents different phase of a flapping cycle. (G,H) Spanwise strains along the veins, each line is normalised by the mean of the strain along the vein at that time. The x-axis (path length, or position along vein) for the spanwise strains are normalised with the total path length for the corresponding vein. (I,J) Temporal variation of the mean spanwise strain on the ventral subcosta and dorsal radius. The measured positions of campaniform sensilla are shown in magenta dots (E-H). The vertical variation of the dots is purely for readability.

## Discussion

We have presented the most comprehensive description of the sensory apparatus on the wings of a flying insect. The mechanical state of a ‘rigid’ wing (as an aircraft) can typically be described by its angle of attack, airspeed, and the configuration of its control surfaces. However, the aeroelastic loading state of the flexible, flapping wings in biology can be far more complicated. As the dragonfly wing twists in resposne to loads, the angle of attack varies along the wingspan with time and the local airspeed is not constant. Thus, to encode the state of a flexible wing, detailed monitoring of both strain and flow sensing are likely to be required. Our findings represent a major advance in our understanding of how mechanosensors in animal wings are organised to fulfil this task.

### Metabolic investment in the dragonfly wing sensory system

One of the largest contributions to the metabolic costs of neuronal signalling is the length of an axon (Niven and Laughlin, 2008; Cherniak, C, 1994; Sterling and Laughlin, 2015). Longer axons increase a neuron’s internal volume, increasing the amount of electrolyte movement required to generate an impulse. Maintaining this ionic balance makes up as much as 50% of the brain’s total energy usage (Grenell, 1958), so there are strong selective pressures for minimizing axon length. Similtaneously, axonal routing must account for practical constraints, for example avoiding regions prone to wing damage, as wings frequently experience wear. For dragonflies, wing damage is almost exclusively found on the trailing edge (Rajabi et al., 2017). Our data suggests that sensory neurons innervating the trailing edge are routed through longitudinal veins that are less prone to damage. If trailing edge sensory neurons were routed along the trailing edge itself, common wing wear could eliminate the function of all sensors distal to the damage.

Another key contributor for metabolic costs is the total neuron count, and the dragonfly wing contains a relatively large number of sensory neurons. Even though our full-wing mapping data is collected from one of the smallest dragonfly species, sensor counts are higher than equivalents reported on the wings of moths, flies or beetles described so far (Dickerson et al., 2014; Cole and Palka, 1982; Frantsevich et al., 2014). Direct comparisons between sensor counts of other insects in the literature are difficult because current reports tend to be incomplete, focusing on specific sensor types rather than a complete sensory ensemble. Each sensor has a cost in terms of mass, moment of inertia, metabolic energy, and increased complexity for decoding. For this reason, we expect biological sensor arrays will use the sparsest sensor arrangements that provide sufficient performance and adequate robustness. In general, dragonfly wing sensors are more sparse posteriorly and distally. Simulations of sparse strain sensor systems show that performance is strongly dependent on placement, with optimised sensor arrays resulting in a significant reduction in sensor count (Mohren et al., 2018). Mechanical stress on wings is largest close to the wing base (Spinola and Chapman, 1975), and the metabolic cost of neuronal signalling increases in proportion to the length of an axon; therefore, a bias towards medial sensor placement likely increases sensory system efficiency. Given that axon diameter tends to be larger for sensors closer to the wing tip for the purpose of conductance speed, the metabolic cost of distal sensors is increased even further.

### Sensory hairs and flow sensing

Here we presume that all innvervated hairs or bristles on the wing are flow sensors. Both the leading and trailing edges are densely populated with sensory bristles. These are easily the most abundant, accounting for approximately 60% of all sensors on the wing. During flapping flight, dragonflies generate a strong leading-edge vortex which is shed during the wing stroke cycle (Thomas, 2004; Bomphrey et al., 2016), and the wing edges tend to experience large fluctuating pressure gradients (Shumway et al., 2020) with cyclic changes of airflow direction. The bristles on the wing margin may, therefore, play a role in detecting the timing and intensity of vortex formation and shedding that could be a useful heuristic for the aerodynamic state. Similar sensors exist in moths and are known to be sensitive to directional vibratory airflow (Rajabi and Gorb, 2020). In the dragonfly, sensory bristles are absent in the proximal 2/3rds of the forewing trailing edge. This area of absence is most likely to be contacted by the hindwing during a flapping cycle. This pattern has been observed in the wing margin bristles of a butterfly’s forewing and hindwing (Yoshida et al., 2001), suggesting a possible common strategy to avoid unwanted signals or damage caused by wing collisions. Apart from these general trends, the wing sensor distribution can be quite diverse across Odonata families. This is likely shaped by the differences in external stimuli (for example, flow velocity field, or dynamic strain field) because of differences in flight kinematics or wing mechanical properties (Combes and Daniel, 2003; Bomphrey et al., 2016; Rajabi and Gorb, 2020).

At the wing’s leading edge, we suspect doubly innervated bristles might serve as a chemosensor as seen in flies (Houot et al., 2017) as well as a mechanosensor (Figure 3A,B). Dragonflies are often considered anosmic due to their minuscule antennal lobes (Fabian et al., 2020), but there is some behavioural evidence suggesting odours are used to choose predation sites (Piersanti et al., 2014). Dragonflies are not able to groom their wings like a fly and their wings do not come into contact at rest, but we cannot exclude the potential tactile functions for wing margin bristles. Other longitudinal veins have isolated bristles with lengths ranging from 20 μm (Figure 3E) to more than 200 μm (Figure 3F), each innervated by a single sensory neuron at its base. The longer bristles have wider bases (3.3-3.4 μm), and resemble the trichoid sensilla found in locusts that detect wind direction and elicit compensatory reflexes (Camhi, 1969; Bacon and Tyrer, 1979; Pflüger and Wolf, 2013; McCorkell, 2016).

To our knowledge, the bristle-bump sensory complex is a novel structure. The bump itself is not innervated by a sensory neuron, and thus plays no direct sensory role. However, it might have a conditioning effect on the external flow stimulus acting on the bristle, altering the tuning properties of the sensor (unpublished data). The bump might also play a protective role for the small bristles, functioning like a roll cage, preventing damage from collisions or parasites. We see evidence that bristle-bumps exhibit isochronicity: the axon diameter varies with axon length, normalising the conduction time of impulses so that all sensors report disturbances to the ganglia with similar latencies. Isochronicity has been reported in motor systems where co-contraction of all motor units are required (Seidl, 2014; Lorenzo et al., 1990). This approach has also been observed in retinal ganglion cells and crayfish antennule (Stanford, 1987; Mellon and Christison-Lagay, 2008). In the example of retinal ganglion cells, the computation of motion vision requires accurate spatiotemporal representation of neighbouring receptive fields. The bristle-bump complex therefore may be monitoring precise spatiotemporal variations in air velocity on the wing surface.

Flow sensor types are clearly associated with local wing morphology, especially corrugation; for example, long trichoid sensilla are almost exclusively found in the wing’s local valleys, while short bristles and bristle-bumps lie on local ridges. This arrangement is suitable to detect the small eddies found within the corrugations as well as the attachment state of airflow (Bomphrey et al., 2016). Bristles without any bump dominate the central part of the wing blade in both the dragonfly and the damselfly. The length of these bristles varies ten-fold, with bristle length decreasing from the wing base to the wing tip. Since the air speed increases from wing root to wing tip during flapping flight, this length distribution may match to the natural patterns of boundary layer height. However it is important to note that flow fields around flapping wings is highly unsteady and may only intermittently follow this pattern. A more detailed aerodynamic study considering the distribution and morphology of the bristles is necessary.

### Strain sensing & possible signal encoding

The isolated campaniform sensilla are sparsely distributed around the edge of the wing. This ring-like arrangement is consistent with capturing the twist of the wing blade efficiently, as the largest deformations occur at a distance from the axis of twist (Mohren et al., 2018; Koehler et al., 2012). Sporadic campaniform sensilla are found along the costa, yet the most posterior campaniform sensilla are located some distance away from the actual trailing edge. This spacing may prevent sensor damage from wing wear, and avoid noise from aeroelastic flutter. Campaniform sensilla counts on the wing of the grasshopper *Melanoplus sanguinipes* (sensilla count: 54, Albert et al., 1976) and the fly *Drosophila melanogaster* (sensilla count: 62, Cole and Palka, 1982; Dickinson and Palka, 1987) are both higher than the odontate OLS regression predicts for wings of those sizes. In general, this suggests that biomechanics and phylogenetic constraints determine sensor counts and larger wings are not simply a scaled-up version of smaller wings. The number of sensors and their specific spacing likely depend on specific details of wing structure and function.

The radius and axillary complex at the wing base are adorned with five fields of campaniform sensilla with 30–90 sensors per cluster (Figure 4A, inset). Both structures are load-bearing, transmitting forces from thoracic muscles to the wings and flight forces from the wings to the body. Severing either structure led to catastrophic failure of wing function (unpublished lab observations). Their sensilla have high aspect ratios (elongated oval shape instead of circular, unpublished data), indicating high directional selectivity (Delcomyn, 1991). Sensor orientations vary within each field, which might reflect loading patterns in the wing base during flapping and gliding flight. Similar fields of campaniform sensilla are found on both the wings and the halteres of flies (Cole and Palka, 1982). These sensors might contribute to inertial sensing or monitoring of wing loading.

Given the structure and material properties of a wing, the strain distribution along the major veins is determined by the sum of inertial and aerodynamic loads. During flapping, inertial loads generally exceed the aerodynamic loads for large insects (Jankauski et al., 2017; Combes, 2003). To encode any aerodynamic components of strain, the wing sensory system must detect small variations of the strain field from the dominant inertial load. Understanding this requires examining wing states and their associated strain distributions on veins. As the first step toward this, we performed structural analysis on the most anatomically accurate dragonfly wing to date. The inertial loading of the wing shows direct strain field propagation along the wing veins with highly stereotyped profiles. Therefore the strain profile on a vein is almost constant in shape (as prescribed by the wing structure and kinematics) but varies in magnitude over time. Any deviation from this profile suggests disturbances due to either aerodynamic events or motor pattern modulation.

How might the wing sensory system encode deformation? Our structural analysis shows that the wing state can be broken down into two components: 1) the temporal profile of strain magnitude over a flapping cycle (Figure 8I,J); and 2) the nominal spatial profile of strain along each vein (Figure 8G,H). Given a specific set of wing-base kinematics, or flapping mode, the spatial strain profile is stereotyped. Therefore, we expect only a few campaniform sensilla are required to encode different flapping modes and the magnitude of the profile. However, any deviation from the nominal spatial strain profile provides a signature for the wing deformation state that could be caused by perturbations or altered kinematics (e.g. during manoeuvring). To detect such variation requires higher spatial resolution and dynamic range in strain sensitivity, which can be accomplished by increasing the campaniform count. Specifically, since campaniform sensilla function as strain detectors, there are two approaches to encode high-resolution strain values. Firstly, clusters of campaniforms at key locations with different strain thresholds and temporal tuning can, collectively, cover a range of strain magnitudes (Delcomyn, 1991). Alternatively, a population of similar campaniforms distributed along the vein can encode the magnitude of variation in their activation phase as the deformation propogates. This is more analogous to a visual system with relatively homogenous photoreceptors which detects motion via phase correlation (Borst et al., 2010). Both approaches require a high campaniform count for high resolution, and are not mutually exclusive. While the observed distribution of campaniform is well-matched to the general strain magnitude, electrophysiological studies are needed to determine the encoding approaches. Finally, strain sensing is only half of the story for wing state estimation. Future fluid-structure interaction simulations will illuminate how flow sensing is tuned to the spatial and temporal aerodynamic state of insect wings.

### Summary and future work

This work provides the most complete map of the wing sensory system for a flying animal, from neuron wiring to sensor morphology and distribution. In addition to the comprehensive maps of one dragonfly and one damselfly species, we compared the sensor distribution on the major veins of the 15 most studied Odonata species. We have described novel sensors and the cross-species comparison has revealed what appear to be conserved sensor placement themes. Finally, our initial, geometrically precise, structural analysis generated specific hypotheses of how the distributed strain sensors monitor wing state. The Odonata wing sensory system is poised to provide information on wing states in addition to providing inertial state estimation for the body. Our findings have raised several key questions regarding aeroelastic sensing for flight control that will be investigated with neurophysiological experimentation and computational fluid-structure interaction modelling.

In soft robotics terms, insect wings are underactuated (controlled degrees of freedom are fewer than the passive degrees of freedom) yet extensively sensorised structures. Their venation patterns must not only produce predictable passive mechanical behaviours, but also provide an appropriate substrate to support the number, density, and appropriate positioning of wing sensors to observe the relevant biomechanical events during flight. This integration of mechanical behaviour and sensing for state estimation (sometimes termed ‘morphological computing’) makes insect wings an intriguing and tractable model for investigating the co-evolution of form, function, and the nervous system.

## Materials and Methods

### Insect specimens

All Odonata specimens were collected in northern Virginia, USA (*), south-east England, UK (**), or obtained from the collection at the Natural History Museum in London, UK. They belong to two suborders: first, Anisoptera, the dragonflies, of which there are 13 families; members of the nine most populous/largest were studied here: 1. Aeshnidae, *Anax junius* (Drury, 1773)*, 2. Chlorogomphidae, *Chlorogompus atkinsonii* (Fraser, 1925), 3. Cordulegastridae, *Anotogaster seiboldii* (Selys, 1854), 4. Corduliidae, *Hemicordulia tau* (Selys, 1871), 5. Gomphidae, *Hagenius brevistylus* (Selys, 1854) and *Stylyrus plagiatus* (Selys, 1854)*, 6. Macromiidae, *Macromia indica* (Frasner, 1924), 7. Libellulidae, *Pantala flavescens* (Fabricius, 1798)*, *Perithemis tenera* (Say, 1840)*and *Sympetrum striolatum* (Charpentier, 1840)**, 8. Petaluridae, *Petalura gigantea* (Leach, 1815), 9. Synthemistidae, *Idionyx stevensi* (Fraser, 1924). Second, Zygoptera, the damselflies, of which there are 35 families. The damselflies used here were from the 1. Calopterigidae, *Calopteryx maculata* (Palisot de Beauvois, 1807)* and 2. Coenagrionidae, *Argia apicalis* (Say, 1840)* and *Ischnura verticalis* (Say, 1839)*. We used males in all our analyses, except for *P. tenera*, for which we used both males and females. In this species, the male wings are ~10% shorter and have ~20% more cross-veins than female wings. However, since most wing sensors are located on the longitudinal veins, the sexual dimorphism is not included in the discussion.

### Sample preparation for confocal imaging

Neurobiotin diffuses at a rate of ~1 cm/24 hr in axons of the diameter found in the Odonata wing, and insect tissue deterioration begins at approximately 48 hr postmortem. Thus, we focused our efforts on one of the smallest Anisoptera, *P. tenera*, and an Isoptera, *A. apicalis*, with comparable wingspan.

Insects were anesthetized on ice and fixed right side up on Sylgard-filled Petri dishes using pins. The right pterothoracic pleural wall and the head were removed, anterior nerves (nerves 1C and 2C in Simmons, 1977) were isolated, placed in a drop (~ 0.7 μl) of distilled water inside a petroleum jelly bowl, and cut. Water was wicked away, replaced with ~0.5 μl of the tracer solution (2 % w/v neurobiotin in dH_2_O, Vector Labs, SP-1120) and the well was sealed closed with petroleum jelly. Insects were kept refrigerated in a humid chamber for 48 hours to allow diffusion of the tracer. The wings were removed by cutting along the basal hinge and fixed in 2% PFA in phosphate-buffered saline with 0.1% Triton X-100 (PBS-T) overnight at 4°C with mild agitation, washed with PBS-T and bleached in 20% peroxide in PBS-T for 48 hr to remove most of the pigment from the cuticle.

When wings from young adults (few hours post-eclosion) were used, oxygen from bleaching formed within the bilayer of cuticle making up the wing blade. This can be explained by the presence of metabolites between the unfused sheets of exoskeleton reacting to the peroxide. At the end of bleaching cycle (indicated by the cuticle’s pale golden hues) the wings became fully inflated by the oxygen. The wings were briefly rinsed in copious amount of water, and placed inside a vacuum desiccator in a PBS-T-filled Petri dish. The low vacuum caused further delamination of the cuticular bilayer. The gas escaped via the openings at the wing base or, occasionally raptured the trailing edge seam. The wings were cut chordwise in half, post-fixed in 2 % PFA for 3 h @ RT, washed and stained with DyLight 594-NeutrAvidin (1:250, Thermo Scientific #22842). The staining reagent now had free/unencumbered access to the neurobiotin-labelled neurons, in PBS with 3% normal goat serum, 1% triton X-100, 0.5% DMSO @ RT with agitation for 2 days. Following washing with PBS-T, the wings were cut near their base and mounted in Tris-buffered (50 mM, pH 8.0) 80% glycerol with 0.5 % DMSO between two coverslips using 350 μm spacers. The wing bases were dehydrated in glycerol (5-80 %), then ethanol (25-100 %), cleared and mounted in methyl salicylate, following modified protocol from Ott (2008).

For treatments of old wing specimens, samples were bleached and re-hydrated using alkaline peroxide (25% H_2_O_2_, 0.2% KOH in water) for 24-48 h and mounted in Tris-buffered (50 mM, pH 8.0) 80% glycerol with 0.5 % DMSO between two 60 mm-long coverslips. The mounting followed the same procedure as fresh wings.

### Imaging and sensor distribution/placement mapping

All samples were imaged on Zeiss 880 upright confocal microscope. Serial optical sections were obtained at 7 μm with a FLUAR 5x/0.25 NA, 2.5 μm with a Plan-Apochromat 10x/0.45 NA objective, at 1.5 μm with a LD-LCI 25x/0.8 NA objective or at 0.5 μm with a Plan-Apochromat 40x/0.8 NA objective. Cuticle autofluorescence and DyLight 594-NeutrAvidin were imaged using 488 and 594 nm laser lines, respectively. The volumes obtained with the FLUAR 5x objective and tiling the wings were stitched in Fiji (Schindelin et al., 2012) (http://fiji.sc/). Images were processed in Fiji, Icy (http://icy.bioimageanalysis.org/) and Photoshop (Adobe Systems Inc.).

The costa, radius and subcosta veins were scanned dorsally and ventrally with Plan-Apochromat 10x/0.45 NA objective (field of view 1.2 × 1.2 mm). To minimize collection time while maintaining high signal to noise ratio, the green autofluorescence of cuticle was excited with 405 and 488 nm lasers at maximum power and the images were collected using single-swipe and minimum pixel dwell time. The vertical aspect of each volume was adjusted to accommodate the imaged vein while the horizontal was kept constant at 1500 pixels. Maximum intensity projections were stitched manually in Photoshop and annotated in Adobe Illustrator (Adobe Systems Inc.) while referring to the volumes viewed in Fiji. 31 sensors were psuedorandomly selected on the forewing and hindwing of one sample for axon path length measurements. Sensors selection was to ensure measurements were taken from a wide variety of wing veins and locations, to avoid wing margin sensors dominating our dataset. Axon path lengths were measured in Fiji, and the minimal path length was determined using the guess and check approach. We constrained our analysis to only allow alternative axon paths that pass through innervated veins.

### Assessment of backfill efficacy

To assess the completeness of neurobiotin labelling, we focused on the hind wing trailing edge bristles, as they are innervated by the longest axons (most challenging) and hence good indicators of labelling efficacy. We found that within 4.4mm stretch the sensory neurons innervating the bristles of the proximal half of trailing edge (i.e., axons passing through anal and cubitus posterior veins along with their tributaries) were all labelled (fig S1 D, top). However, within the distal 5.8 mm stretch of the trailing edge (i.e. axons traveling through media, radius veins and associated cross-veins), 2/23 dorsal and 4/33 ventral sensory neurons were not labelled (89.3 % efficacy) (Fig S1 D, bottom). Projecting from the full labelling of the proximal half, we accounted for this ~10% dropout at the distal portion.

### Cell body volume and axon diameter quantification

For the cell body and axon diameter quantification, equally distributed wing sensors were selected from the right hindwing of one *P. tenera* sample. Cell body volumes were quantified by first manually identifying the intensity threshold in ITK SNAP (Yushkevich et al., 2006) (http://www.itksnap.org/) which was used to segment the cell body volume voxels from the background image and axon. Second, the identified threshold of 1800 was implemented in a custom written Fiji macro to automatically segment all images and obtain the resulting cell body volume. The results were validated with the manually obtained volumes from ITK SNAP for randomised samples. Axon diameter was measured at 20 μm proximally from each cell body in Fiji.

Assuming a cylindrical axon with homogenous membrane properties, the neural conduction speed *v* can be estimated as (Tasaki, 2004):

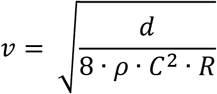

where *d* is the axon diameter, *ρ* is the resistivity of the axoplasm, *C* is the membrane capacitance per unit area, and *R* is the resistance per unit area. The conduction latency *t*, therefore, can be represented as:

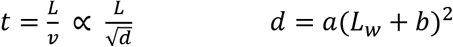

where *L*_w_ is the axon length in the wing, a is a scaling factor and b is the axon length offset accounting for the distance from the wingbase to the thoracic gangalion. We performed nonlinear least squares fit to this relationship with all the data and with different combinations of sensor types.

### Wing sensor distribution quantifications

Based on the annotations, sensors were counted and their positions digitized in Fiji. The sensor density was defined as the inverse of the averaged sensor spacing over two consecutive sensors (sensors/mm). The sensor position was normalised by the spanwise wing length for comparison across species. Sensor density was smoothed by a moving average over 7 data points and subsequently normalised by the mean sensor density of the respective sensor type on the vein to facilitate relative comparisons across species.

The relationship between sensor count and wing length for each sensor type was analysed via an ordinary least squares (OLS) regression in Rstudio (Rstudio Team, 2020). The lower and upper confidence intervals (2.5% and 97.5%) of the OLS regression were calculated. In this analysis, the sensor count for each sensor type corresponded to the sum of the sensors of the first three anterior veins (costa, subcosta and radius).

### Wing geometry modelling for structural analyses

A male *Sympetrum striolatum* dragonfly was euthanised in a freezer overnight. The entire body was stained by elemental iodine, which also had a desiccating effect (Boyde et al., 2014). The x-ray microtomography (μCT) scanning on the left hindwing was performed using a SkyScan 1172 scanner (Bruker, Belgium). The wing was mounted with the long axis vertical, aligned to the rotational axis of the scanner, and five sub-scans were performed. In total, 12445 images were taken at pixel size of 2.83 μm and exposure time of 2.6 s, with a rotation step of 0.08° for 180° rotation for each sub-scan. The source voltage and current were 65 kV and 153 μA, respectively. The raw images were processed in Bruker NRecon to obtain cross-sectional slices. The images were then imported to ORS Dragonfly v3.6 (Object Research Systems, Canada) for registration and three-dimensional reconstruction. A mesh file (PLY) was extracted by choosing an appropriate intensity threshold. After cleaning and reduction in MeshLab, the mesh was imported to Rhinoceros 6 (Robert McNeel & Associates, USA). The vein network was approximated with circular cross-section pipes whose diameter and joint positions were informed from the mesh. The membrane was generated for each cell surrounded by veins with uniform thickness of 10 μm, which was determined by sampling the measured cross section using the Dragonfly software. The veins and membranes were imported into AutoDesk Inventor Professional 2020 (Autodesk, Inc., USA) to merge them into a single body. The wing model was exported in STEP format and imported into ANSYS Mechanical Application 2019 R3 (ANSYS, Inc., USA), where the mesh for finite element analysis was generated for transient structural simulations.

### Wing dynamic loading computational solid dynamics (CSD)

The transient structural simulation was performed with ANSYS Mechanical Application 2019 R3. The wing model mesh consists of quadratic tetrahedron elements. The uniform material property was assumed for simplicity, where density was 1200 kg m_−3_, Poisson ratio was 0.3, and the Young’s modulus was 5 GPa. These values were derived from previous work [S6], where the Young’s moduli were 6 GPa for vein and 3.75 GPa for membrane. Single-axis rotation around the wingbase resembling flapping is the simplest and most natural dynamic loading stimulus. Thus only 1 degree of freedom for rotation around the flapping axis (Figure 5A–D) was prescribed, and other 5 (3 translational and 2 rotational) degrees of freedom were fixed. The flapping motion is described as:

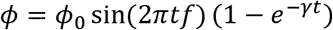

where *ϕ* is positional angle given at the wingbase in degrees (Figure S3D, blue line), *ϕ*_0_ is the desired wingbeat semi-amplitude and set to 30°, *t* is time in seconds, *f* is wingbeat frequency and set to 40 Hz, and *γ* is a factor to change how fast the amplitude approaches to *ϕ*_0_. The wingbeat amplitude increases gradually in this formulation so we can avoid the abrupt start and it is expected to arrive at the periodic deformation state earlier than a simple sinusoidal flapping. The factor γ was set to 30, which results in the wingbeat amplitude is 30% of *ϕ*_0_ for the 1st cycle but 67%, 84%, and 93% for 2nd, 3rd, and 4th cycles, respectively. At 10th cycle the amplitude is 99.9% of *ϕ*_0_. The wing started flapping from the mid-downstroke phase at time *t* = 0, and the computation was performed until time *t* = 16*T*= 0.4 s, where *T* = 1/*f* was the wingbeat period (i.e. 0.025 s). The time step size (time increment, *Δt*) for each computational step was determined to be 3×10^−4^ s after several preliminary trials. No external forces such as gravitational force or aerodynamic force were considered in this analysis.

The positional angle was defined such that the original attitude at mid-downstroke is 0, and negative angle at supination and positive angle at pronation. The wingtip path and wing positional angle at the wingtip were calculated for troubleshooting.

### Mesh convergence check

Coarse and fine meshes were generated for mesh convergence verification, where coarse mesh has 845,039 nodes while the fine mesh has 1,226,124 nodes (Figure S3A,B). They showed slightly different low-frequency oscillation patterns in the wingtip positional angles (Figure S3C–H). For fair comparison, we took the wingbeat cycles where the mean positional angle during downstroke is the smallest for each case (25th cycle for coarse mesh and 30th cycle for fine mesh, figure S3G,H). The spanwise strains in both meshes showed good agreement (Figure S3I). Thus we can confirm simulation convergence and show the results from fine mesh alone.

### Strain analysis

The resultant strain field was evaluated in two ways: the normal elastic strain along two major longitudinal veins (ventral side of subcosta and dorsal side of radius), and the normal elastic strain contours on the wing surface at four time instants (pronation, mid-downstroke, supination, and mid-upstroke). Here, the normal elastic strain was computed for the wing radial direction for each instant by coordinate transformation using the instantaneous positional angle. We will refer this strain component as “spanwise strain” for short.

The period of low-frequency oscillation for the fine mesh case was around 8-10 wingbeat cycles (Figure S3D,F,H). The strain along two veins for one low-frequency period (from the 22nd to 30th wingbeat cycles) showed some variation but the general trend was consistent, where the mean spanwise strain for the ventral subcosta and the dorsal radius show opposing temporal patterns (Figure S3J,K,L). Therefore, in the main text, we show the representative results from the 28th wingbeat cycle starting from pronation (i.e., 27.75≤*t*/*T*≤28.75). For simplicity, each wingbeat cycle was described with 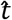 such that 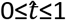, where 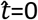 and 1 are pronation and 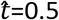 is supination. The spanwise strain along two veins were analysed in three ways. The raw strain along each vein for each instant was plotted (Figure 5E,F); the raw strain was normalised with the mean along the vein at the corresponding time (Figure 5G,H); and the mean strain was plotted against time. This way, we expected to see if the time-varying strain distribution can be decomposed into spatial and temporal components.

The spanwise strain contours on the surface were calculated in ANSYS CFD-Post 2019 R3. The mesh and strain for each time instance was exported in a Generic format using Command Editor with simple for-loop. The files and converted into VTK (legacy) format with a custom-written Python script and imported into ParaView 5.9.0 (KitWare, Inc., USA) for visualisation.

### Modelling limitations

To our knowledge, this wing model is the most geometrically detailed insect wing to date, yet the model and the simulation both have some limitations. The material properties (in particular the Young’s modulus) is uniform over the entire wing; no elastic joints (e.g. nodus) are implemented yet; cross-sectional shapes of the veins are circular and filled (in reality, it is hollow and more elliptical in dorsoventral direction). Furthermore, we subjected the wing to simple sinusoidal curve around a single flapping axis, whereas active rotation in the other two axes occurs in real flapping flight. Nevertheless, this current model illustrates general strain propagation patterns necessary to infer how aeroelastic wing state might be encoded. We will continue to improve the model fidelity in future work.

## Author contributions

HTL & RJB conceived the project and secured funding; JF & HTL performed pilot studies and preliminary mapping; IS performed confocal microscopy, and generated the neuroanatomy maps; IS and MU analysed the sensor distribution data; MM built the wing model and performed the dynamic loading simulation. JF & HTL developed the manuscript; all authors revised the manuscript for submission.

## Acknowledgements

We would like to thank the Biotechnology and Biological Sciences Research Council for supporting this work (Grant BB/R002509/1 to HTL and BB/R002657/1 to RJB). We would like to thank Dr Ben Price for lending us selected specimens from the NHM collection. We thank Mr Mark Hopkinson and Skeletal Biology Group in the Royal Veterinary College for their help in the μCT scanning. We would like to thank Dr Maria Jose Freeman for assistance with the acquisition and maintenance of Odonata specimens. We would like to thank Dr Alexandra Yarger and Dr David Labonte for their valuable feedback on the manuscript. We lament the environmental destruction for housing development at one of our dragonfly sites in SE England.

---

**Supplemental Figure 1.**
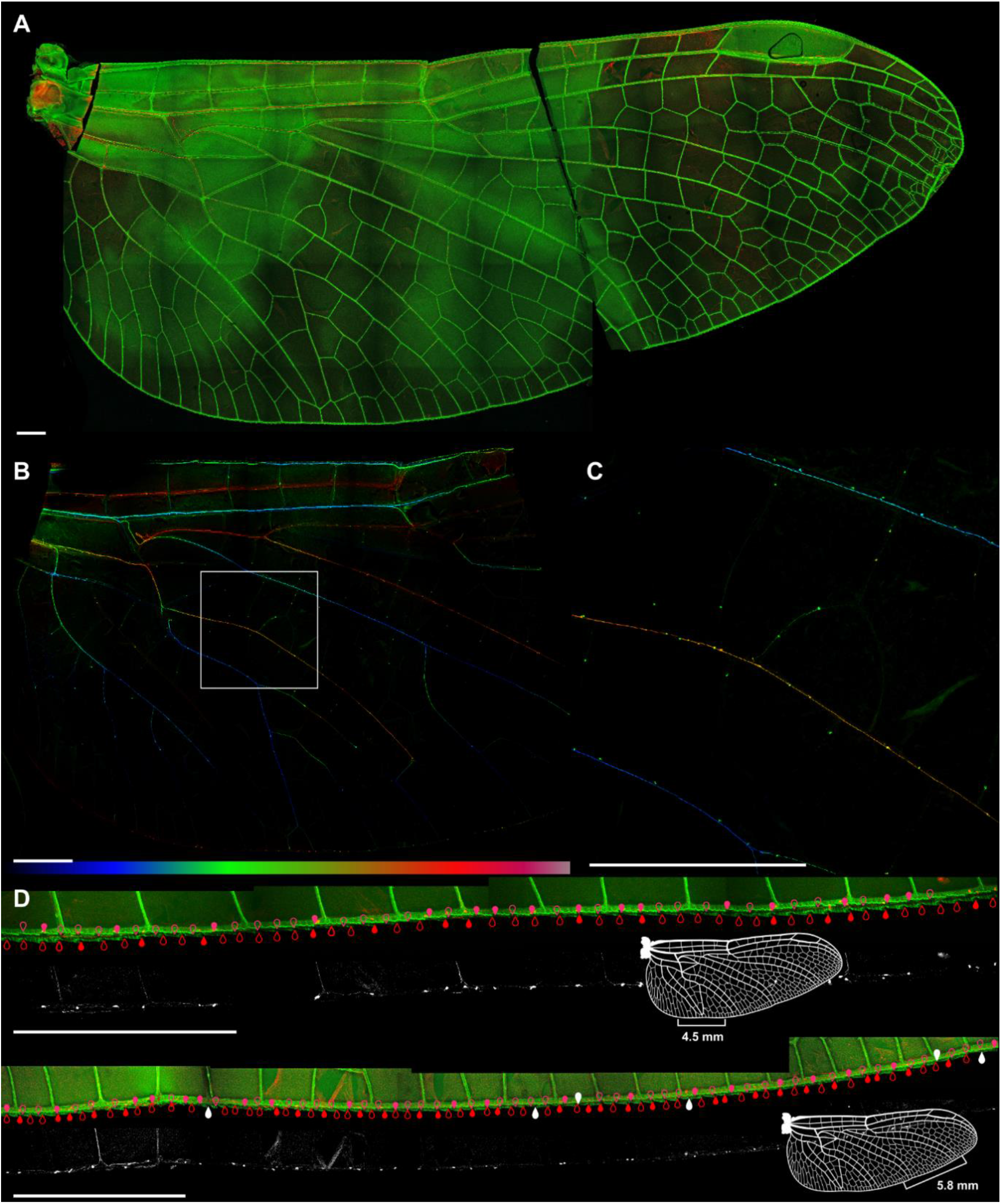
**(A)** *Perithemis tenera* hind wing; maximum intensity projection of stitched volumes collected with the FLUAR 5x objective. Cuticle autofluorescence – green; DyLight 594-neutravidin – red. **(B)** Depth color-coded projection of the red channel. Soma of various sensory neurons appear as dots. **(C)** Area indicated in (A) by the box. **(D)** Assessment of the efficiency of the backfill using *P. tenera* hind wing trailing edge bristle-bump complexes as a proxy. Filled markers indicate labelled cell bodies; empty markers – bumps not accompanied by a bristle; white markers: unlabeled cell bodies (bristle present, lack of detectable signal from the cell body). Scale bars: 1 mm.

**Supplemental Figure 2.**
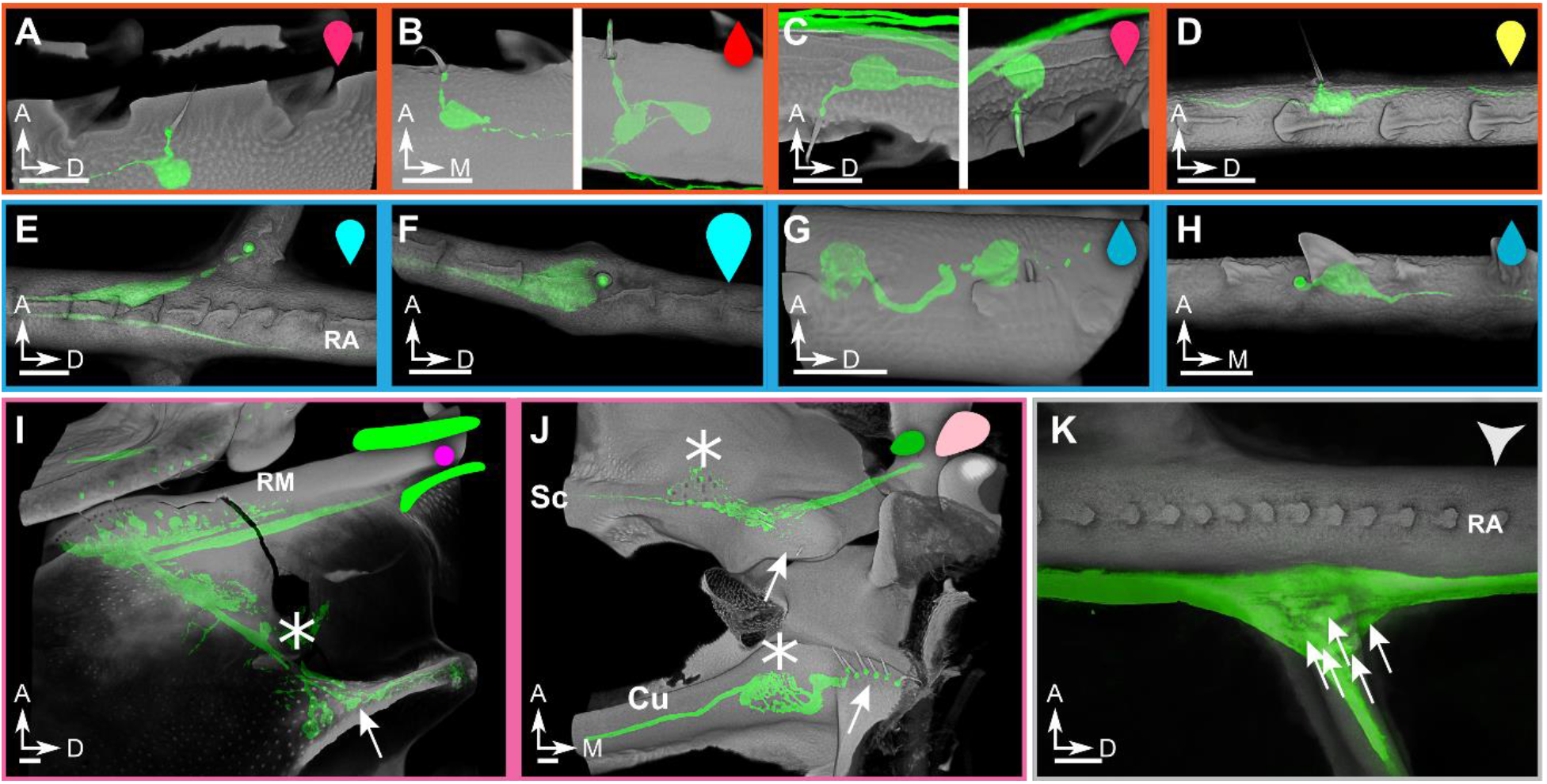
Examples of wing veins and base sensors of damselfly *Argia apicalis*. Most structures are analogous to the dragonfly. **(A-D)** air flow sensors; **(A)** dorsal costa bristle-bump complex; **(B)** ventral costa bristle-bump complex; left: single-innervated; right: double-innervated bristle. **(C)** trailing edge bristle-bump complex; left: single-innervated; right: double-innervated bristle. **(D)** short isolated bristle. **(E-H):** strain sensors; (**E**) Campaniform sensilla (CS) of a cross vein medial to pterostigma; (**F**) large dorsal radius anterior CS immediately distal to pterostigma; **(G)** ventral media CS proximal to the wing base; (**H**) one of ventral media posterior CS. **(I-K)** wing base sensors. **(I)** crevice organ (i.e. campaniform sensilla) – two parallel fields of directionally tuned/elongated CS dorsally at the base of radius/media (RM) vein. Asterisk marks the dorsal insertion site of a wing base chordotonal organ. Arrow – CS field of axillary complex posterior ridge. **(J)** Ventral view of the subcostal and cubitus vein base. Arrows point to hair plates, asterisks mark CS fields. **(K)** Multipolar receptor at the junction of radius and a cross vein posterior to the nodus. Arrows point to several large cell bodies. All the images show mechanosensors of the right forewing. Cuticle’s autofluorescence shown in grey, neurons labelled with Neurobiotin/DyLight-594-Neutravidin – in green. Scale bars: 25 μm. The symbols in the upper right corner of each panel correspond to the markers showing placement of sensors in Fig. 3.

**Supplemental Table 1.** Mean sensor density, wing length and sensor count of 15 Odonata species. Mean sensor density and sensor counts are presented for the three major longitudinal veins Costa (C), Subcosta (Sc) and Radius (R) on dorsal (d) and ventral (v) wing side. Mean sensor density is presented for campaniform sensilla (cs), isolated bristles (is), bristle-bump complexes (bb) and small bristles in the range of a bump (sis). The last sensor type was assigned to the bristle-bump complexes in the article because the bristles had the same morphology and could be associated to a nearby cuticular bump.

|  | Mean sensor density (1/mm) |  |  |  |  |  |  |  |  |  |  |  | Wing length<br>(mm) | Sensor<br>count |
| --- | --- | --- | --- | --- | --- | --- | --- | --- | --- | --- | --- | --- | --- | --- |
|  | C(d) |  | Sc(d) | R(d) |  |  | C(v) |  | Sc(v) |  |  | R(v) |  |  |
|  | cs | bb | is | cs | bb | sis | cs | bb | cs | bb | sis | is |  |  |
| *I.verticalis* |  | 6.69 |  |  |  |  |  | 9.64 |  |  |  |  | 1.15 | 202 |
| *P.tenera* |  | 7.81 | 5.58 | 2.2 | 2.92 |  |  | 11.98 | 10.73 |  |  | 3.71 | 2.07 | 519 |
| *A.apicalis* |  | 2.95 |  |  |  |  |  | 5.94 |  |  |  |  | 2.46 | 168 |
| *S.striolatum* | 1.45 | 4.99 | 3.78 | 1.06 | 2.65 |  |  | 7.83 |  | 2.19 |  | 3.45 | 2.95 | 547 |
| *C.maculata* |  | 4.93 |  |  |  | 6.71 | 2.27 | 8.86 |  |  |  |  | 3.06 | 392 |
| *I.stevensi* | 1.94 | 10.63 | 3.75 | 1.33 | 2.48 |  |  | 15.57 | 6.71 | 3.88 |  | 11.19 | 3.31 | 1080 |
| *H.tau* | 2.38 | 5.5 | 7.24 | 1.45 | 2.7 |  |  | 9.12 | 4.46 | 2.39 |  | 12.16 | 3.63 | 734 |
| *S.plagiatus* | 3.08 | 4.01 |  | 5.52 | 1.5 |  | 2.10 | 6.21 | 7.18 | 1.45 |  |  | 3.74 | 600 |
| *C.atkinsoni* | 2.55 | 5.18 |  | 6.96 | 2.48 |  | 4.25 | 8.74 | 6.69 |  |  | 12.64 | 4.05 | 900 |
| *P.flavescens* | 1.16 | 4.94 | 12.37 | 1.06 | 2.37 |  |  | 7.73 |  | 2.63 |  | 14.41 | 4.33 | 844 |
| *M.indica* | 1.11 | 6.72 |  | 2.86 | 3.27 |  |  | 15.18 | 3.37 | 3.05 |  | 7.3 | 4.76 | 1420 |
| *A.junius* |  | 5.47 | 2.99 | 2.35 | 2.24 |  | 2.83 | 6.47 | 5.08 | 2.2 |  | 7.55 | 5.36 | 1065 |
| *P.gigantea* | 2.98 | 4.22 |  | 5.36 | 2.00 |  | 3.58 | 4.02 | 7.38 | 1.43 | 0.98 | 4.45 | 5.41 | 793 |
| *A.sieboldii* | 2.34 | 3.93 | 1.64 | 5.49 | 1.99 | 3.73 | 1.75 | 7.26 | 6.99 |  | 1.64 |  | 5.44 | 958 |
| *H.brevistylus* | 1.77 | 3.93 | 2.03 | 3.98 | 1.81 | 2.62 | 2.63 | 6.55 | 5.81 |  | 2.03 |  | 5.45 | 766 |

**Supplemental Figure 3.**
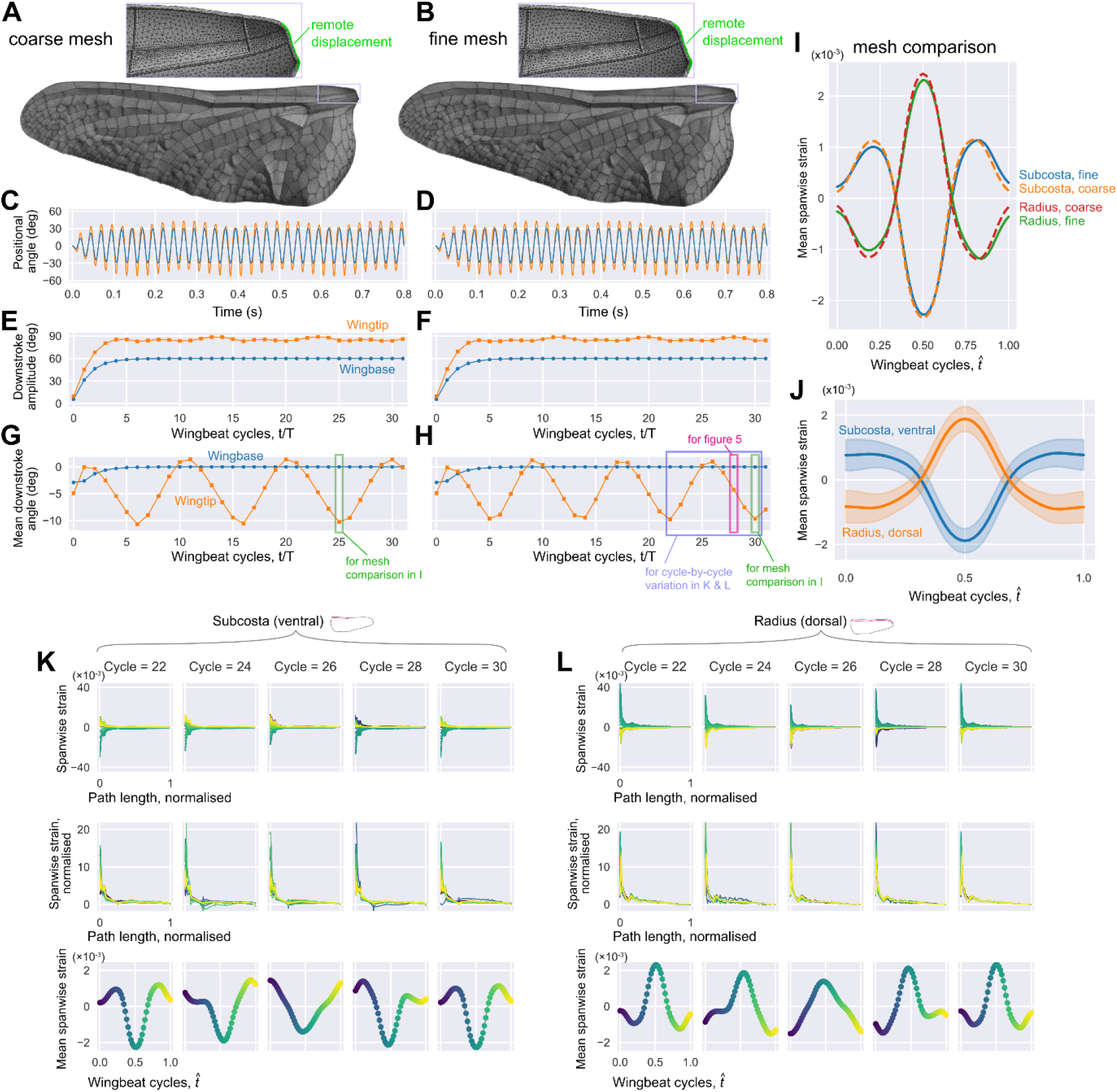
Mesh convergence and cycle-by-cycle variation in spanwise strain. Coarse **(A)** and fine **(B)** meshes. Enlarged views show the mesh near the wing base. The green region is the faces that the “remote displacement” boundary condition was applied in ANSYS Mechanical to drive the wings. Positional angles **(C,D)**, wingbeat amplitude for downstroke **(E,F)**, and mean positional angle for downstroke **(G,H)**, where blue lines are for wingbase and orange lines are for wingtip. Note the 0 mean positional angle means the wing flaps equal amount for dorsal and ventral directions. In the current case, the mean positional angle tends to be negative, indicating the downward bending. The mean (along each of 2 vein) spanwise strains for coarse and fine meshes **(I)**, where blue and green solid lines are subcosta and radius in fine mesh, respectively, and orange and red dashed lines are subcosta and radius in coarse mesh, respectively. The ensemble average for the mean spanwise strain over nine consecutive wingbeats (21.75 ≤ *t/T* ≤ 30.75) (J), where blue line is subcosta and orange line is radius. Mean and ±1 SD are shown. The cycle-by-cycle variation in spanwise strain for subcosta **(K)** and radius **(L)**. Cycle 22 corresponds to 21.75 ≤ *t/T* ≤ 22.75, and so on.

